# Drug repurposing to face Covid-19: Celastrol, a potential leading drug capable of inhibiting SARS-CoV-2 replication and induced inflammation

**DOI:** 10.1101/2021.04.20.439992

**Authors:** Carlos A. Fuzo, Ronaldo B. Martins, Thais F.C. Fraga-Silva, Martin K. Amstalden, Thais Canassa De Leo, Juliano P. Souza, Thais M. Lima, Lucia H. Faccioli, Suzelei C. França, Vania L.D. Bonato, Eurico Arruda, Marcelo Dias-Baruffi

**Affiliations:** Departamento de Análises Clínicas, Toxicológicas e Bromatológicas. Faculdade de Ciências Farmacêuticas de Ribeirão Preto, Universidade de São Paulo, Ribeirão Preto, SP, Brazil; Departamento de Biologia Celular e Molecular e Bioagentes Patogênicos, Faculdade de Medicina de Ribeirão Preto, Universidade de São Paulo, Ribeirão Preto, SP, Brazil; Departamento de Bioquímica e Imunologia. Faculdade de Medicina de Ribeirão Preto, Universidade de São Paulo, Ribeirão Preto, SP, Brazil; Unidade de Biotecnologia, Universidade de Ribeirão Preto, Ribeirão Preto, SP, Brazil

**Keywords:** Covid-19, SARS-CoV-2, drug repurposing, reverse gene signature, molecular docking, celastrol

## Abstract

The global emergence of Covid-19 has caused huge human casualties. Clinical manifestations of the disease vary from asymptomatic to lethal, and the symptomatic form can be associated with cytokine storm and non-homeostatic inflammatory response. In face of the urgent demand for effective drugs to treat Covid-19, we have searched for candidate compounds using a drug repurposing approach based on *in silico* analysis followed by biological validation. Here we identified celastrol, a pentacyclic triterpene isolated from *Tripterygium wilfordii* Hook F – a plant used in traditional Chinese medicine – as one of the best compounds out of 39 repurposed drug candidates. Celastrol reverted gene expression signature from SARS-CoV-2-infected cells; bound with high-affinity energy to viral molecular targets such as main protease (M^pro^) and receptor-biding domain (RBD); inhibited SARS-CoV-2 replication in monkey (Vero and Vero-ACE2) and human (Caco-2 and Calu-3) cell lines; and decreased interleukin-6 (IL-6) secretion in SARS-CoV-2-infected human cell lines. Interestingly, celastrol acted in a concentration-dependent manner, with undetectable signs of cytotoxicity. Therefore, celastrol is a promising lead drug candidate to treat Covid-19 due to its ability to suppress SARS-CoV-2 replication and IL-6 production in infected cells, two critical events in the pathophysiology of this disease.

## 1. Introduction

Covid-19 (Coronavirus Disease 2019), caused by the β-coronavirus SARS-CoV-2 (Severe Acute Respiratory Syndrome Coronavirus 2), was first reported to the World Health Organization in January 2020 after a local pneumonia outbreak of unknown etiology in Wuhan, China^1,2^. SARS-CoV-2 has been rapidly and effectively transmitted from human-to-human and became a worldwide pandemic that affected more than 137 million people in April 2021^3,4^.

The devastating effects of Covid-19 on global public health and economy have demanded urgent efforts to discover potential drug and vaccine candidates to prevent and treat this disease. Many immunization strategies have been studied since the beginning of the pandemic, demonstrating the fast and extraordinary achievement of pharmaceutical companies in the development of vaccines for Covid-19. Although there are 289 vaccine candidates under development, from which 20 were under phase 3 clinical trial in February 2021^5–7^, there are many challenges to achieve an efficient global immunization, such as production limitations, efficacy levels, restrictions on use, dosing procedures, storage requirements, price, emergence of SARS-CoV-2 mutants, and promotion of durable immunological memory^7–12^.

Despite the availability of effective vaccines, the quick discovery of drugs to prevent and treat SARS-CoV-2 infection is a critical demand to face Covid-19^13^. The drug reuse strategy accelerates the discovery of candidate compounds with known activities associated with reducing the SARS-CoV-2 viral load or promoting a better clinical evolution of Covid-19^14,15^. One aspect of the immunopathology of this disease that stands out is non-hemostatic inflammation associated with cytokine storm involving by several mediators, including interleukin-6 (IL-6), which is a severity biomarker of this illness^16–18^.

In this sense, bioinformatics and computational biology are powerful and multidimensional *in silico* tools for drug discovery and repurposing of compounds approved or under clinical trial^19–22^. Some of the currently employed strategies to find drugs applicable in Covid-19 have focused on *(i)* host or virus targets, such as the receptor-binding domain (RBD) present in spike glycoprotein and angiotensin-converting enzyme II (ACE2), which mediate virus-host cell interaction^23,24^; *(ii)* proteins/enzymes from virus biosynthesis machinery, such as main protease (M^pro^) and RNA-dependent RNA polymerase (RdRp)^25–27^; and *(iii)* reversion of the host gene expression signature of SARS-CoV-2-infected cells^15,28,29^. This study used *in silico* predictions to repurpose drug candidates that could concomitantly reverse the SARS-CoV-2 gene expression induced in host cells, including IL-6, and target proteins/enzymes essential to the SARS-CoV-2 life cycle, followed by biological validation of the best candidate drug repurposed.

## 2. Models and Methods

### 2.1. Identification of signatures from SARS-CoV-2 *in vitro* infection model and search for drugs with reversed viral infection signature

The gene expression signature from SARS-CoV-2-infected primary human lung epithelium cell line (NHBE) was obtained from Blanco-Melo and co-authors^30^, by filtering differentially expressed genes (DEGs) after differential expression analysis from independent biological triplicates of SARS-CoV-2 (USA-WA1/2020 strain)-infected and mock-treated cells. The signature was constructed based on DEGs with absolute value of fold change in log_2_ scale greater than 1 (|log_2_(FC)| > 1), and significance accepted at adjusted *p*-value (*p*_adj_) smaller than 0.05 (*p*_adj_ < 0.05), as determined by the Benjamini and Hochberg (BH) method^31^.

The obtained signature containing up- and downregulated genes was used to search for drugs with reversed gene expression signature in comparison to viral infection. The best-ranked drugs (*Q*_score_) from the signatures of small molecule expression profiles in LINCS L1000 dataset ^32^ from Characteristic Direction Signatures Search Engine (L1000CDS^2^) were listed^33^ (https://amp.pharm.mssm.edu/L1000CDS2/). The signature was submitted to over-representation analysis using Reactome pathways^34^ and *clusterProfiler R* package^35^ to find biological pathways that were enriched due to viral infection, within BH adjusted *p*-value < 0.05. The DEGs from enriched pathways were used to construct a biological score (*B*_score_) for each drug ranging from 0 to 1, where 0 indicated no reversion and 1 indicated total reversion of the main enriched pathways. A weighting factor was calculated for each DEG by multiplying their occurrence by |log_2_(FC)|. *B*_score_ was the sum of DEG weighting factors and it was normalized by the sum of all weighting factors. The graphical representation of drug signatures data and scores were generated with *pheatmap*^36^ and *ggplot2* packages^37^ in *R* software^38^.

### 2.2. Docking of selected drugs on SARS-CoV-2 protein targets

The compounds capable of reversing SARS-CoV-2 infection signature (selected in Section 2.1) were submitted to molecular docking on three SARS-CoV-2 protein targets: the catalytic site of M^pro^, the RNA binding site of RdRp, and the RBD domain of spike glycoprotein, using multiple structural conformations. Here we used ensemble docking to increase the sampling and avoid unique conformational bias. We have previously generated the representative structures for RBD (unpublished results) using ten centroid clusters obtained from a 600 ns trajectory, with the aid of molecular dynamics simulation based on the deposited co-crystal RDB/ACE2 structure (PDB id 6M0J)^39^. The multiple structures of M^pro^ and RdRp were obtained from *Protein Data Bank (PDB)*^40^ (Table S1). The heteroatoms were removed from M^pro^ and RdRp structures, the resulting structures were repaired, and the energy was minimized with *FoldX*^41^. *AutoDockTools*^42^ was used to prepare drug and protein input structures for docking analysis and grid box definitions (Table S1). Docking analysis was carried out using *Autodock Vina*^43^ with exhaustiveness parameter equal to 20. Graphical representations of energy results were plotted with *ggplot2* R package and molecular model structures were drawn with *Pymol*^44^ and *Discovery Studio Visualizer^®^ (version-2020)*^45^.

### 2.3. The rationale for selecting a predictable candidate lead drug for experimental validation

*In silico* analysis guided the selection of a candidate drug for biological validation, considering the drug's ability to reverse the genetic signature of SARS-CoV-2 infection and its binding affinities to molecular viral targets. Briefly, the ten best repurposed drugs were selected based on *Q*_*score*_ values. A compound with high predicted median binding affinity energy for RBD, M^Pro^, and RdRp – and thereby with strong potential to inhibit the functions of these molecular targets – was selected using molecular docking data.

### 2.4. Preparation of viral stock

To obtain SARS-CoV-2 viral stock, clinical isolates (SARS-CoV-2 Brazil/SPBR-02/2020 strain) from RT-PCR-confirmed Covid-19 patients were propagated using monkey Vero CCL-81 cells (kidney), under strict biosafety level 3 (BSL3) conditions. Briefly, for initial viral passages, Vero CCL-81 cells were cultured in Dulbecco minimal essential medium (DMEM) supplemented with 10% heat-inactivated fetal bovine serum (FBS) and antibiotic/antimycotic mix (10,000 U/mL penicillin and 10,000 μg/mL streptomycin). Viral inoculum (1:100 ratio) was added to the cells, and the culture was incubated (48 hours, 37 °C, 5% CO_2_ humidified atmosphere) in DMEM without FBS but supplemented with antibiotic/antimycotic mix and trypsin-protease inhibitor, L-1-tosylamide-2-phenylethylchloromethyl ketone (TPCK) host cell treatment (1 μg/μL) to optimize virus adsorption to the cells ^46^. After confirming the cytopathic effects of the viral preparation using an inverted Olympus ix51 microscope, infected Vero CCL-81 cells were detached by scraping, harvested, and centrifuged (10000 ×*g*, 10 minutes, room temperature). The resulting supernatants were stored at −80 °C until use. Finally, virus titration was performed on Vero CCL81 cells using standard limiting dilution to determine the 50% tissue culture infectious dose (TCID50) of viral stock^47,48^.

### 2.5. *In vitro* SARS-CoV-2 infection

SARS-CoV-2 infection was assessed *in vitro* in four cell lines: Vero CCL-81, human ACE2-transfected Vero CCL-81 (Vero CCL-81-ACE2), human Calu-3 (lung), and human Caco-2 (colon). Cells were seeded into 24-well plates (80,000 cells/well) to ensure 90% of confluence on the day of inoculation/infection. The four cell lines were infected with SARS-CoV-2 and treated with celastrol. Cells were infected with SARS-CoV-2 at multiplicity of infection (MOI) 1.0 in 500 µL of infection media composed of DMEM without FBS, 1% antibiotic/antimycotic mix, and 1 μg/μL trypsin-TPCK. After 2 hours of incubation, supernatant containing SARS-CoV-2 was removed and replaced by celastrol (125, 250, 500, and 1,000 nM) or vehicle (0.05% DMSO) diluted in fresh medium, followed by 48 hours of incubation at 37 °C and under 5% CO_2_. Photomicrographs were taken using the Olympus ix51 inverted microscope and analyzed using the QCapture Pro 6.0 software under 200× magnification (QImaging)^49^, in order to examine whether celastrol interfered with SARS-CoV-2 cytopathic effects in Vero CCL-81 cells. The supernatants were collected for RNA extraction and viral load was quantified using a standard curve. All the infections were conducted in technical triplicate.

### 2.6. Cell viability

Cytotoxicity of celastrol (Sigma-Aldrich) to Vero CCL-81, Vero CCL-81-ACE2, Calu-3, and Caco-2 was determined using the Alamar Blue Cell Viability protocol (Thermo Scientific, Waltham, USA), according to the manufacturer's instructions. Briefly, the cells were seeded into a 96-well plate to grow as monolayers and treated with celastrol (250 or 1000 nM) in DMSO (0.05%) or DMSO (5%, v/v; cell death reference) for 24 hours. Alamar Blue reagent (10% v/v) was added to the cells, and the plate was incubated at 37 °C, for 4 hours. Median fluorescence intensity was measured using the SpectraMax i-3 (Molecular Devices) microplate reader, with excitation and emission wavelengths set at 530 and 590 nm, respectively. The mean value from the control (untreated cells) was set as 100%, and the viability of cells from each treatment condition was calculated relative to this value, in triplicate.

### 2.7. RT-PCR for SARS-CoV-2

The SARS-CoV-2 genome was quantified using primer-probe sets for N2 and RNAse-P housekeeping gene, following USA-CDC protocols (Table 1)^50^. To determine the genome viral load from *in vitro* infection assays, N2 and RNAse-P gene were tested by one-step real-time RT-PCR using total nucleic acids extracted with Trizol^®^ (Invitrogen, CA, EUA) from 250 µL of culture supernatants. All RT-PCR tests were carried out using the Step-One Plus real-time PCR thermocycler (Applied Biosystems, Foster City, CA, USA). Briefly, 100 ng of RNA were used for genome amplification, mixed with specific primers (20 μM), probe (5 μM), and TaqPath 1-Step qRT-PCR Master Mix (Applied Biosystems, Foster City, CA, USA). The following reaction parameters were used: 25 °C for 2 minutes, 50 °C for 15 minutes, 95 °C for 2 minutes, followed by 45 cycles of 95 °C for 3 seconds and 55 °C for 30 seconds.

**Table 1.**
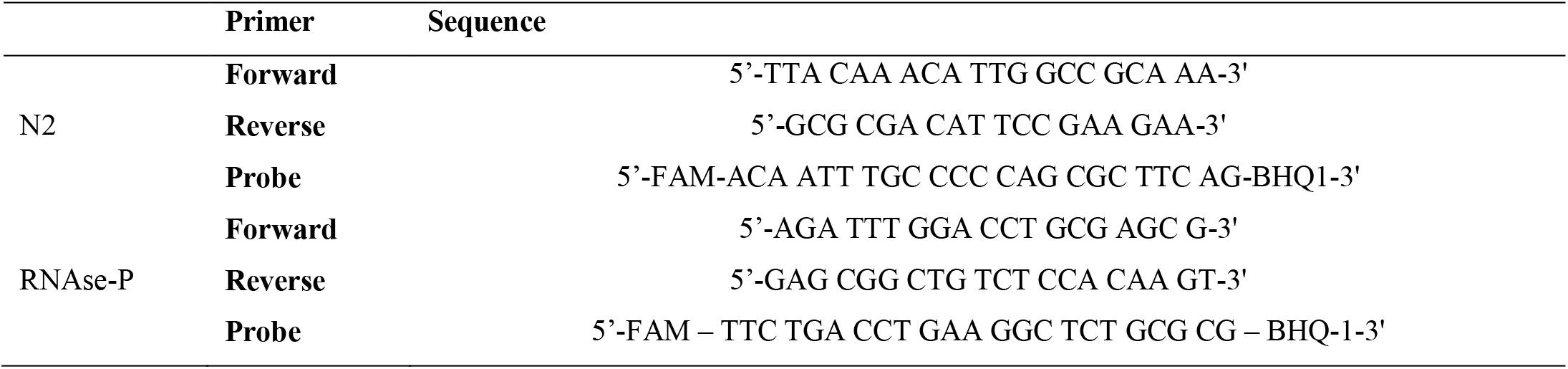
Primers/probe sequences for detection of SARS-CoV-2 genome and housekeeping gene^50^

A plasmid standard curve was plotted to determine SARS-CoV-2 viral load. A 944 bp amplicon was inserted into a TA cloning vector (PTZ57R/T CloneJetTM Cloning Kit Thermo Fisher^®^), starting from residue 14 of N gene, which includes all three sets of primers/probe designed by the CDC protocol (N1, N2, and N3). To quantify the amount of virus produced, a tenfold serial dilution of the plasmid was prepared in the range from 10^6^ to 1 plasmid copy. The coefficient of determination (*R*^2^) for the plasmid standard curve was 0.999, with efficiency above 91% reached using any set of primers/probe^51^.

### 2.8. Interleukin 6 quantification

IL-6 levels were quantified in Caco-2 and Calu-3 cell culture supernatants using the Human DuoSet ELISA assay kit (R&D Systems, USA), according to the manufacturer’s instructions. The IL-6 detection limit was 9.38-1200 pg.mL^−1^.

### 2.9. Statistical analysis

Raw data for viral load from RT-PCR was transformed into log scale, normalized, and analyzed by non-linear regression using dose-response curve fitting and the equation log (inhibitor) *versus* normalized response with variable slope. All the experimental data are expressed as mean ± standard error of the mean (SEM), and they were plotted and analyzed using GraphPad Prism 7 software^52^. Statistical significance was analyzed by one-way analysis of variance (ANOVA) followed by Tukey's post-test for multiple comparisons. Differences were considered significant at *P* < 0.05. The *P* values were labeled as * *P* < 0.05, \*\**P* < 0.01, \*\*\**P* < 0.001, and \*\*\*\**P* < 0.0001.

## 3. Results

### 3.1. Identification of potential candidates for drug repurposing to treat Covid-19

We determined the genetic signature of SARS-CoV-2-infected NHBE cells based on transcriptome data reported in the literature^30^. Then, we used |log2(FC)| > 1 and *p*_adj_ < 0.05 as selection criteria, and identified 64 up- and 8 down-regulated DEGs in infected NHBE cells (Table S2). The over-representation analysis of selected DEGs in *Reactome pathways* annotations indicated the existence of 22 significantly enriched biological pathways (Fig. 1a, Table S3). The top enriched pathways were related to interleukin, chemokine, and interferon signaling, according to literature data^30^, and were closely related to biological pathways involved in SARS-CoV-2 human infection^53–56^.

**Figure 1.**
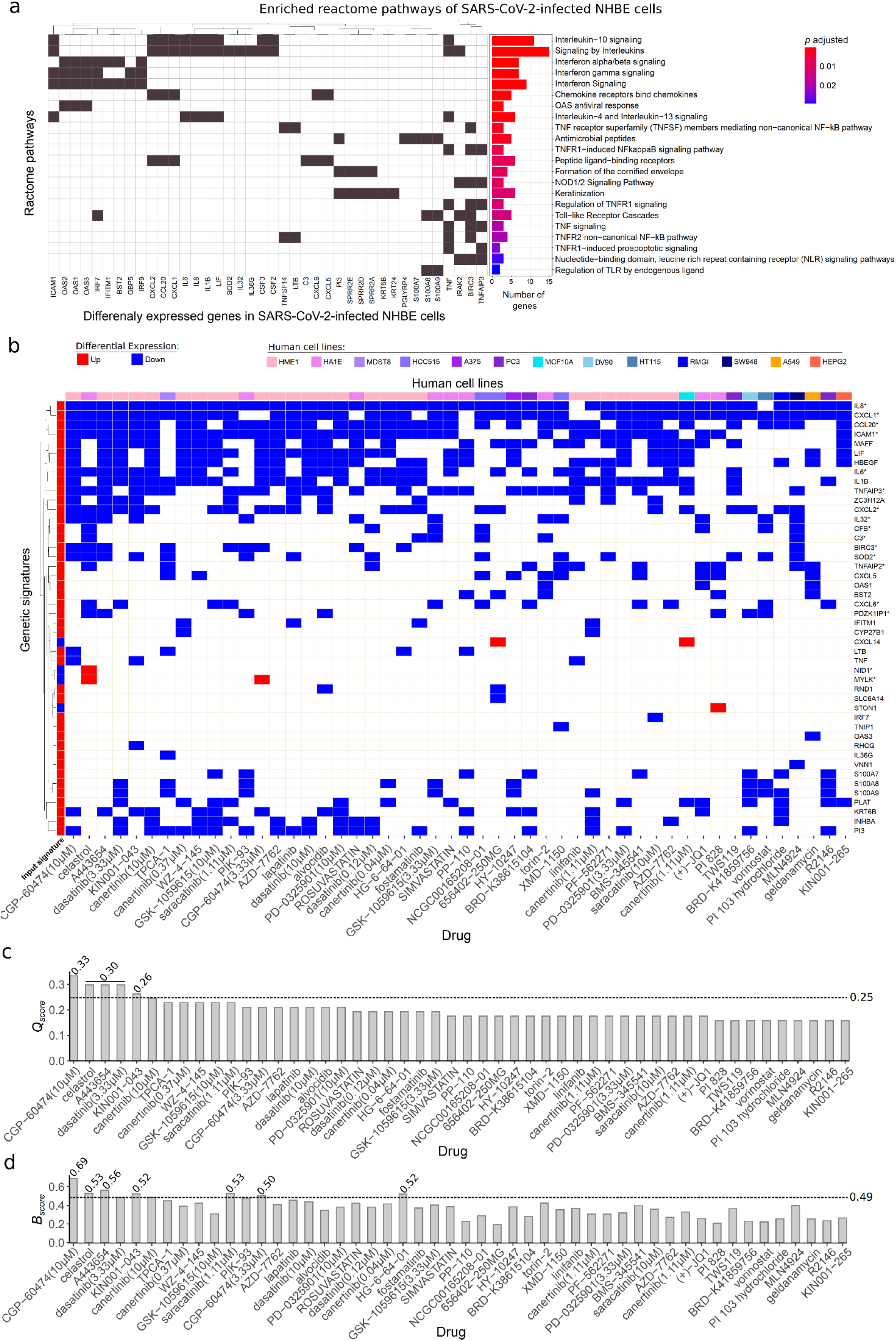
Assessment of gene signature in SARS-CoV-2-infected human bronchial epithelial cells (NHBE) for drug repurposing to treat Covid-19. (**a**) Enriched biological pathways associated with Covid-19 from *Reactome* annotations (*padj* <0.05) from DEGs identified in NHBE-infected cells transcriptome data. The genes that enriched each pathway (left) are indicated together with statistical results and pathway description (right). (**b**) Heatmap of genes from the 50 best-ranked drug signatures that reversed the genetic signature of SARS-CoV-2-infected NHBE cells in decreasing order of *Q*_*score*_. The signature map annotations are related to up- and down-regulated genes, and cell lines are indicated in different colors. (**c**) Drugs in decreasing order of *Q*_*score*_ following the output of L1000CDS2. A dashed line indicates the mean *Q*_*score*_ (0.26) threshold. Equal *Q*score values are displayed over the bars. (**d**) *B*_*score*_ for each drug, considering the enrichment analysis and drug reverse signature. The dashed line corresponds to the mean *B*_*score*_ (0.49). *Genetic signature that justified the biological validation of celastrol.

Considering that the genetic signature described here was composed of DEGs associated with Covid-19 pathophysiology, we searched for candidate drugs that could reverse the genetic signature of NHBE-infected cells using the L1000CDS2 protocol to define the *Q*_*score*_. The top 50 better-scored drug signatures were able to modify the expression pattern of 46 DEGs (42 up-regulated and 4 down-regulated) from the original input of 72 DEGs (Fig. 1b; Table S4). Based on the *Q*_*score*_ values from these 50 drug signatures, a medium value score was defined as 0.26 (Fig. 1c). CGP-60474, a cyclin-dependent kinase inhibitor^57^, exhibited the best-ranked *Q*_*score*_ (0.33): it down-regulated 19 DEGs that were up-regulated by SARS-CoV-2 infection. The second best-ranked *Q*_*score*_ (0.30) was associated with three drugs that modified the expression levels of 17 DEGs induced by SARS-CoV-2: *i*) celastrol, a pentacyclic triterpenoid derived from *Tripterygium wilfordii* Hook F with anti-inflammatory and anti-cancer properties^58^; *ii*) A443654, a potent inhibitor of all members of the protein kinase B (Akt/PKB) family^59^; and *iii*) dasatinib, a small-molecule inhibitor of multiple tyrosine kinases^60^ (Fig.1c). In the set of best-ranked drugs by *Q*_*score*_, CGP-60474, celastrol, and A443654 also presented *B_score_* values higher than the mean score of 0.49 (Fig. 1d), suggesting that they are potential reversers of biological pathways related to SARS-CoV-2 infection.

### 3.2. Selection of drugs that bind to viral molecular targets (RBD, M^pro^, and RdRp) among those that reverse SARS-CoV-2-induced genetic signature

We have searched for candidate drugs with two potential actions: to revert the genetic signature of SARS-CoV-2 infection and interfere with critical events of viral infection. For this purpose, 39 drugs previously identified as reversers of the viral genetic signature were submitted to molecular docking on four regions of three critical viral targets: *i*) the catalytic site of M^pro^; *ii*) two sites of RBD, RBD1 and RBD2; and *iii*) RNA-binding cleft of RdRp (Fig. 2a).

**Figure 2.**
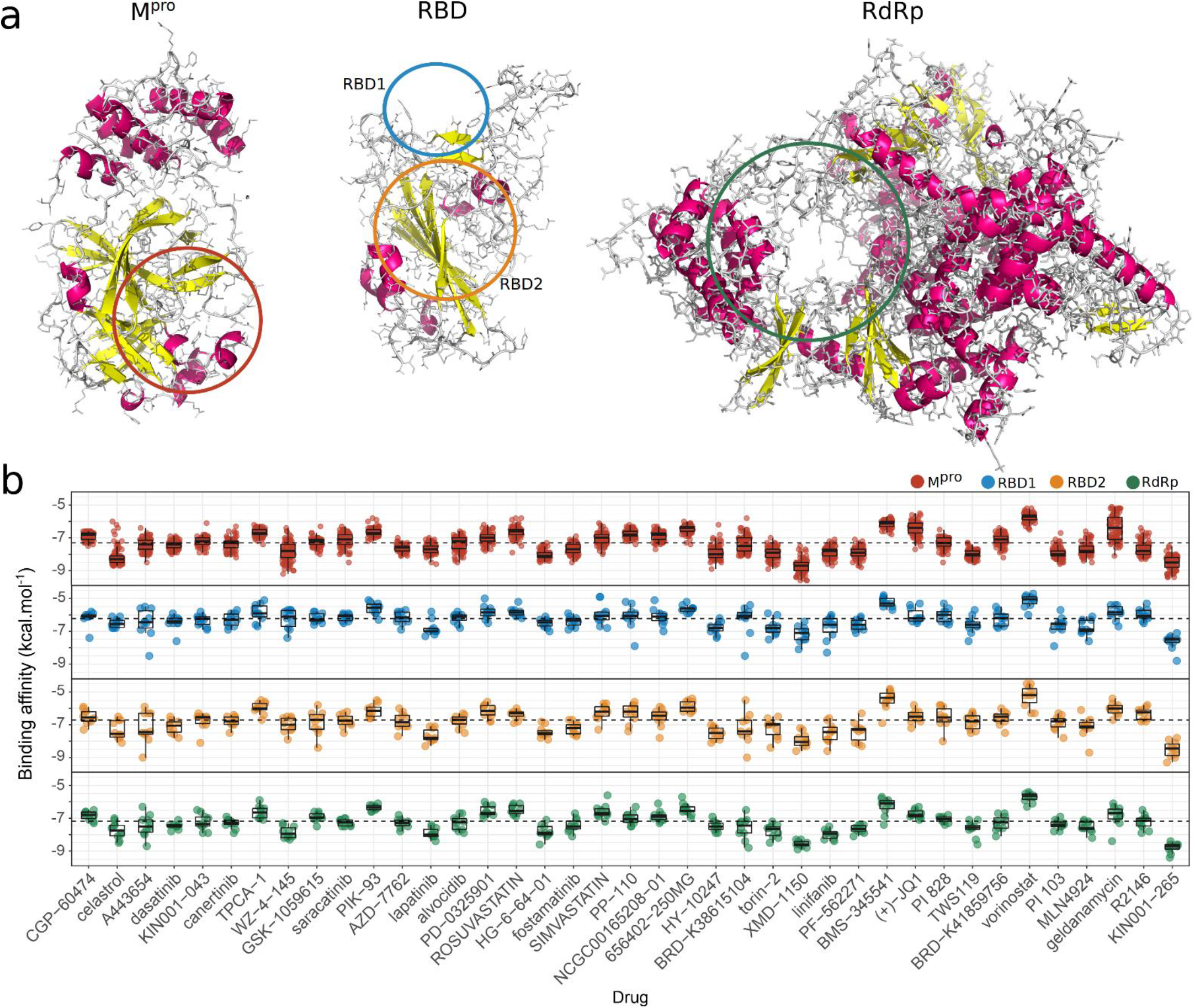
Docking analysis of molecular interactions between viral targets and drugs capable of reversing SARS-CoV-2 genetic signature. **(a)** Representative structures of the SARS-CoV-2 molecular targets (M^pro^, RBD, and RdRp), where atoms are represented as lines and secondary structures as a cartoon with helices highlighted in magenta and sheets in yellow. Colored line circles (red - M^pro^; RDB1 - blue; RBD2 - orange; RdRp - green) indicate the binding sites used for docking related to known inhibition of critical regions from each viral target ^19,24–27^. **(b)** Boxplots illustrate the affinity binding energies (kcal.mol^−1^) obtained from docking analysis between several structural conformations of 39 drugs and each viral site (M^pro^ = 83; RDB1= 10; RBD2 = 10; and RdRp = 10 structures). The reverser drugs were sorted based on decreasing order of *Q*_*score*_ that indicated their potential to revert the genetic signature of SARS-CoV-2 infection. Dotted lines indicate median affinity binding energies defined for each viral target (M^pro^ = −7.3; RDB1= −6.2; RBD2 =−6.7; and RdRp = −7.2 kcal.mol^−1^) considering all investigated drugs.

We determined the pose with the highest binding affinity energy (*ΔG*_*bind*_ in kcal.mol-1) for each ensemble’s structures of all selected compounds and viral molecular targets (Fig. 2b). The *ΔG*_*bind*_ values ranged from −9.6 kcal.mol^−1^ for the leucine-rich repeat kinase inhibitor XMD-1150^61^ binding to M^pro^ to −4.5 kcal.mol^−1^ for the histone deacetylase inhibitor vorinostat^62^ binding to RBD2. The ensemble dockings on different conformations led to energetic variations but not to wide dispersion of *ΔG*_*bind*_ values, demonstrating that certain drugs maintained their high binding affinity to different conformations of the studied targets.

Celastrol exhibited the most attractive *ΔG*_*bind*_ median values of energy out of the first ten best-ranked drugs for the viral targets (M^pro^ = −7.3, RDB1= −6.2, and RBD2 =−6.7 kcal.mol^−1^), and presented the second most attractive *ΔG*_*bind*_ median value for RdRp (−7.8 kcal.mol-1 - Fig. 2b). The chemical pattern of celastrol-target interactions was composed of conventional hydrogen bonds, carbon-hydrogen bonds, Pi interactions, and van der Waals interactions that favored a more attractive binding affinity of celastrol to M^pro^, RBD1, and RBD2 with *ΔG*_*bind*_ equal to −8.7, −6.8, and −8.1 kcal.mol-1, respectively (Fig. 3). The celastrol binding to M^pro^ catalytic site was mediated by the sigma Pi interaction among the E-ring O_2_, His41 and hydrogen bound in Thr25, and OH group of A-ring C3. The mean atomic distance between the B-ring C6 and the Cys145 sulfur atom – a critical amino acid of the M^pro^ catalytic site – was 0.63 nm (range: 0.43-1.33 nm) (Fig. 3), suggesting an appropriate geometry to the formation of a covalent bond. In addition, celastrol exhibited one of the highest *B*_*score*_ values (0.53) among the 39 drugs analyzed, indicating that this compound had a great potential to revert the genetic signature of SARS-CoV-2 infection (Fig. 1d). Of note, the protein kinase inhibitor WZ-4-145^63^ displayed the best *ΔG*_*bind*_ (−8.3 kcal.mol-1) to RdRp due to a myriad combination of interactions, including conventional hydrogen bonds, alkyl, and Pi interactions (Fig. 2b and Fig. 3). Some drugs with high affinity to the studied targets, such as XMD-1150 to M^pro^ and KIN001-265 to RBD1, RBD2, and RdRp, were poorly ranked in reverse signature repurposing but they could be relevant due to their predicted binding abilities. The *ΔG*_*bind*_, *Q*_*Score*_, and *B*_*score*_ values of celastrol supported its selection to biological validation.

**Figure 3.**
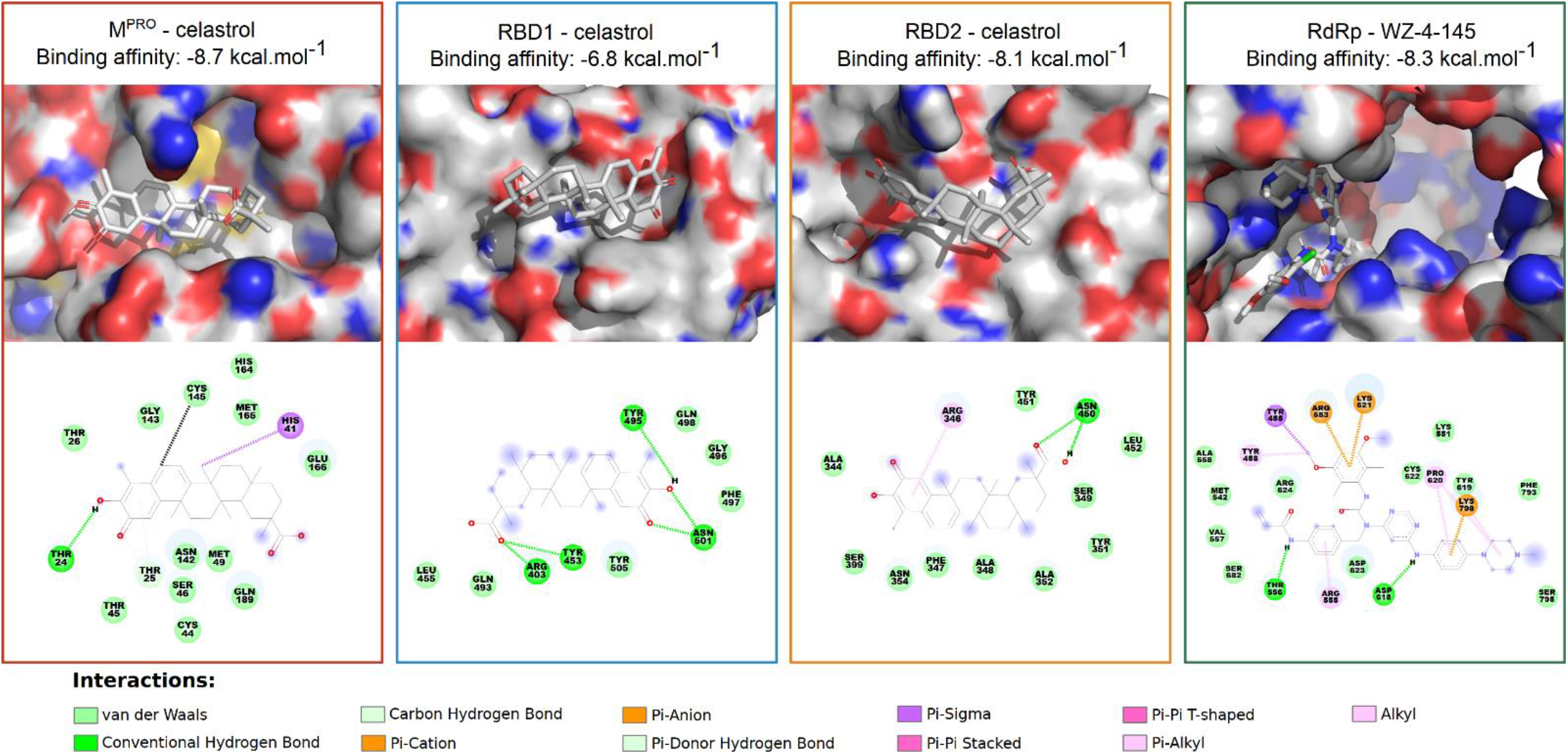
Detailed chemical interactions between the best-ranked drugs and inhibition sites of SARS-CoV-2 molecular targets. 2D target-drug interaction diagrams for the best structural configurations of viral molecular targets was determined using the Discovery Studio^®^ software (version-2020). Celastrol had the most attractive *ΔG*_*bind*_ median values of affinity energy to three targets (Mpro, RBD1, and RBD2) and WZ-4-145 to one (RdRp). The distance between the B-ring C6 and the sulfur atom of the Cys145 residue, which may be related to a possible Michael adduct formation for the best energy poses in each M^pro^ structure, ranged from 0.43 to 1.33 nm, with average value of 0.63 nm (black dashed line).

### 3.3. Celastrol inhibits SARS-CoV-2 cellular propagation and IL-6 secretion by infected cells without cytopathic effect *in vitro*

#### 3.3.1. Celastrol reduces SARS-CoV-2 viral load on infected cells

*In silico* analysis indicated that celastrol is a potential drug candidate to treat Covid-19 due to its ability to reverse the genetic signature of SARS-CoV-2 infection and interact with high affinity with critical molecular viral targets. This *in silico* prediction was validated using *in vitro* SARS-CoV-2 cell propagation inhibition assays. Celastrol significantly reduced viral load in two monkey cell lines (Vero CCL-81 and Vero CCL-81-ACE2) and two human cell lines (Caco-2 and Calu-3) (Figure 4). Interestingly, the anti-SARS-CoV-2 effect of celastrol was concentration-dependent and it drastically inhibited viral replication in Vero CCL-81, Vero CCL-81-ACE2, and Calu-3 cells when tested at 1000 nM.

**Figure 4.**
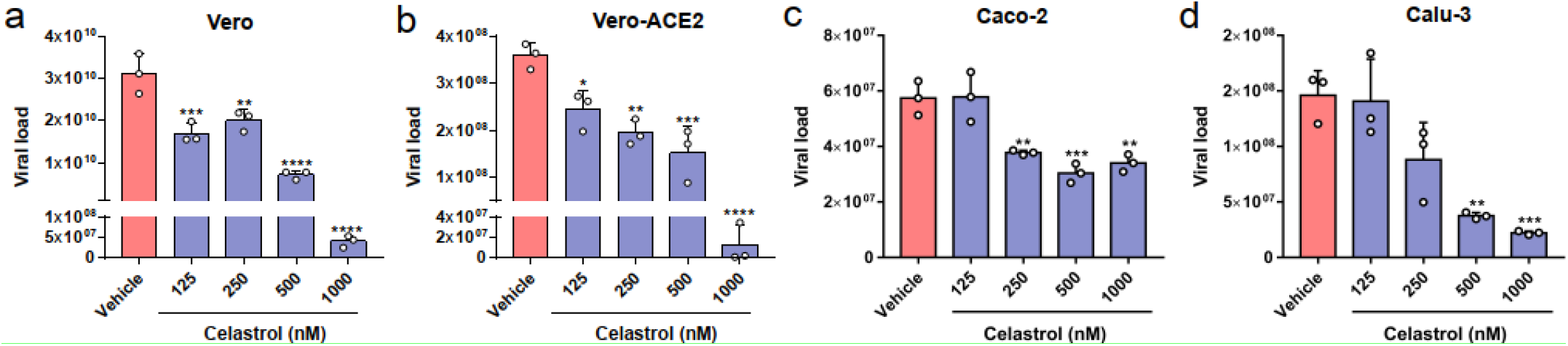
Celastrol suppresses SARS-CoV-2 in vitro propagation in non-human and human cell lines. RT-PCR quantification of SARS-CoV-2 RNA in supernatants from the infected cell lines (**a**) Vero CCL-81 (**b**) Vero CCL-81-ACE2, (**c**) Caco-2, and (**d**) Calu-3 treated with celastrol at concentrations of 125, 250, 500, and 1000 nM. DMSO solution (0.05%; vehicle) was used as negative control. Cells were infected using MOI = 1.0 for 2 hours and then treated with celastrol for 48 hours. The detection levels of SARS-CoV-2 RNA were performed in the supernatants of cultures and expressed in viral load using a standard curve described in the Material and Methods section. Statistical differences between celastrol treatments and the negative control were analyzed by ANOVA followed by Tukey’s post-test. The significance levels were indicated as * P < 0.05, **P < 0.01, ***P < 0.001, and ****P < 0.0001.

To examine whether cytotoxicity mediated the anti-viral effect of celastrol, we determined the cell viability of the four cell lines treated with celastrol at two concentrations, 250 nM and 1000 nM, which promoted its minimal and maximum anti-viral action in human cell lines, respectively. Celastrol-treated cells and the negative control had similar viability levels, of nearly 100% (Fig. 5a-d). Treatment with 5% DMSO (positive control) decreased cell viability by more than 80%. Celastrol also significantly reduced SARS-CoV-2 cytopathic effect in a concentration-dependent manner (Fig. S1).

**Figure 5.**
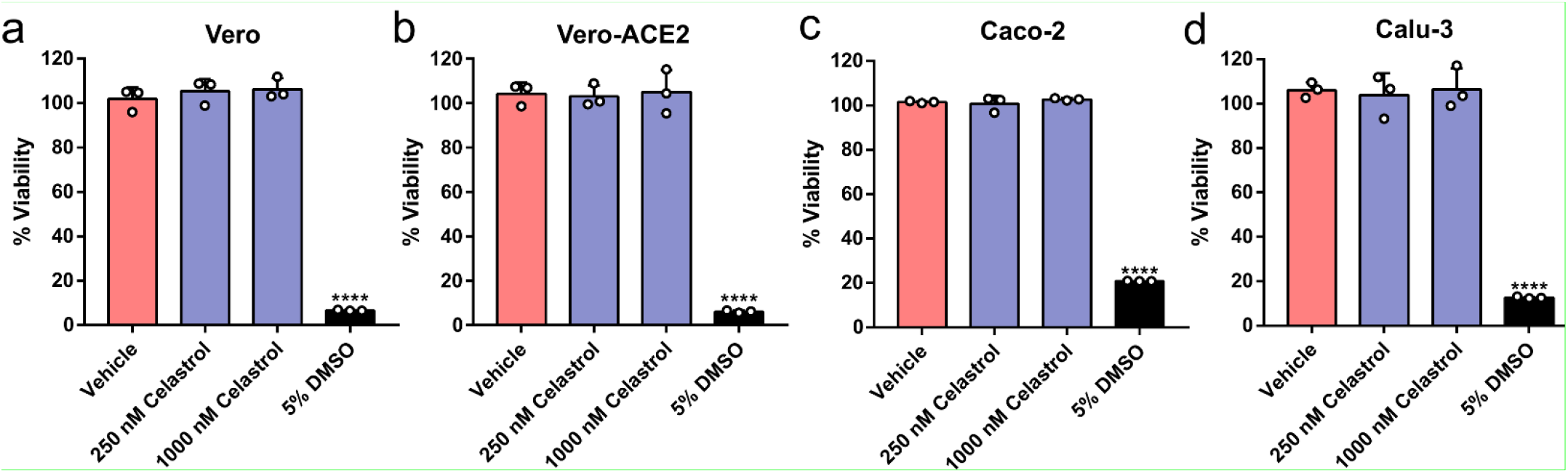
Celastrol does not affect cell viability. AlamarBlue assay was used to measure the potential cytotoxic effect of celastrol. The monkey cell lines (**a**) Vero CCL-81 and (**b**) Vero CCL-81-ACE2, and the human cell lines (**c**) Caco-2 and (**d**) Calu-3 were treated with celastrol at 250 and 1000 nM, for 48 hours. DMSO solutions at 0.05% and 5.0% were used as the negative and positive controls of cell death, respectively. Statistical analysis was performed using ANOVA followed by the Tukey's post-test to compare treatment with celastrol and 5.0% DMSO. ****P < 0.0001.

#### 3.3.2. Celastrol lowers IL-6 production in SARS-CoV-2-infected human cell lines

SARS-CoV-2 infection of human cell lines increased gene expression of inflammatory mediators, such as IL-6, as demonstrated by *in silico* analysis, and celastrol reversed this effect (Fig. 1). To test this prediction, we determined IL-6 levels in the supernatant of SARS-CoV-2-infected cells treated or not with celastrol. This compound at 500 and 1000 nM significantly decreased IL-6 production by Caco-2-infected cells, and at 1000 nM it decreased IL-6 production by Calu-3-infected cells (Fig.6). Curiously, Caco-2-infected cells produced around ten-fold lower IL-6 levels than Calu-3-infected cells. This phenomenon may be associated with the lower viral load in culture supernatants of Caco-2 cells, as compared with samples from Calu-3 cells (Fig. 4c,d).

**Figure 6.**
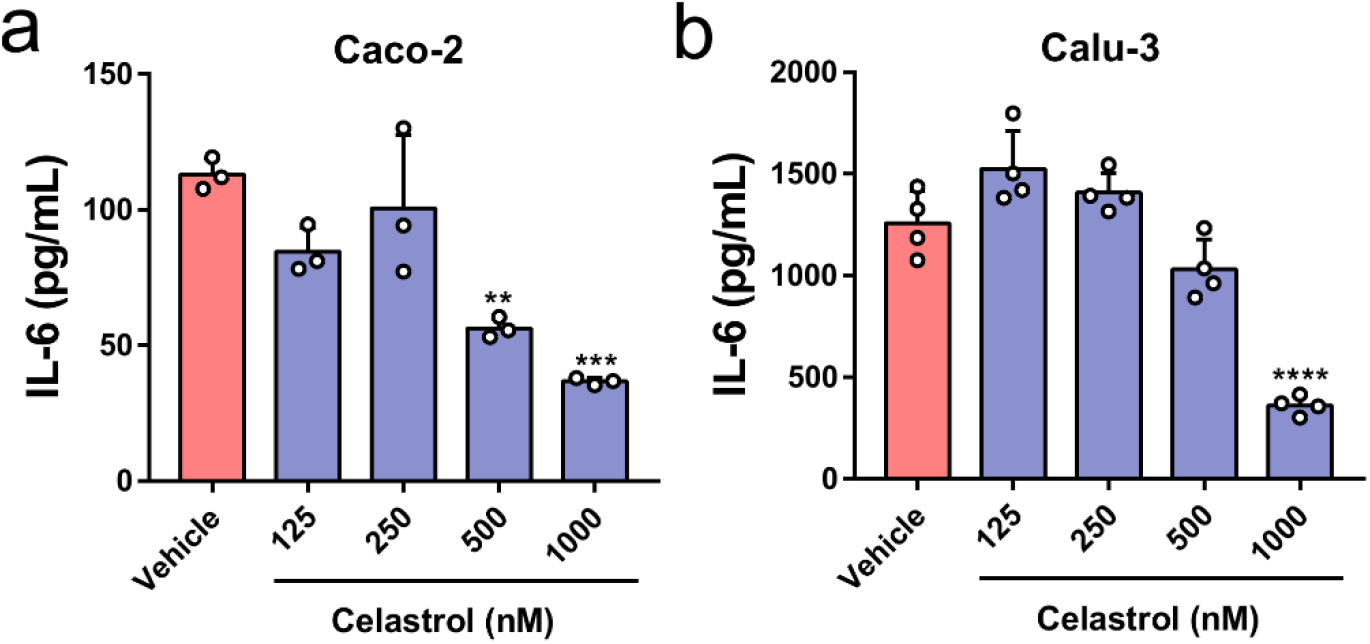
Celastrol inhibits IL-6 production in SARS-CoV-2-infected human cell lines. ELISA quantification of IL-6 in the culture supernatants from the human cell lines (**a**) Caco-2 and (**b**) Calu-3, infected with SARS-CoV-2 (MOI = 1.0) for 2 hours and then treated with celastrol for 48 hours. DMSO solution at 0.05% (vehicle) was used as the negative control. Statistical differences between celastrol treatment and control were analyzed by ANOVA followed by Tukey’s post-test. The significance levels were indicated as **P < 0.01, ***P < 0.001, and ****P < 0.0001. The results are expressed as mean ± standard error of the mean (SEM) of IL-6 (pg/mL) and are representative of two independent experiments with Caco-2 cells and one experiment with Calu-3 cells.

## 4. Discussion

Here, we identified 39 potential repurposed drug candidates for Covid-19 treatment based on *in silico* prediction of their abilities to revert the genetic signature of SARS-CoV-2-infected cells and bind with high affinity to viral targets. One of the *in silico* best-ranked candidates was celastrol, which was also able to inhibit the virion particles release and IL-6 secretion by SARS-CoV-2-infected cells. This combination between computational and experimental analyzes validated the literature predictions about the potential use of celastrol to face Covid-19 based on its anti-inflammatory and antiviral properties^64,65^. Other *in silico* best-ranked candidates with antitumor, kinase-inhibition, and anti-inflammatory properties have been proposed to treat Covid-19, such as CGP-60474, A-443654, alvocidib, canertinib, dasatinib, vorinostat, and geldanamycin^29,66–68^.

The perturbed gene expression during SARS-CoV-2 infection *in vitro* modulates biological processes of clinical relevance, such as cytokine-mediated gene expression, type I interferon pathways, and antiviral responses as detected in our current analysis and literature^30^. As expected, genes of pro-inflammatory cytokines (IL-1β, IL-6, IL-8, TNF-α, and others), chemokines (CXCL1 and CCL20), and some NF-κB pathway components were up-regulated, and are related to the severity and progression of Covid-19^55,69–72^.

Celastrol was the second best-ranked drug able to reverse the genetic signature in SARS-CoV-2-infected cells, in agreement with its reported anti-inflammatory properties^73^. This triterpene also dampens HIV-1 Tat-induced inflammatory responses, inhibits the production of proinflammatory chemokines, such as CXCL10, IL-8, and MCP-1 (YOUN et al., 2014), and suppresses replication of Dengue virus serotype 1-4 by promoting IFN-α expression and stimulating downstream antiviral responses^74^. Celastrol suppresses the production of cytokines during cytokine storm and of chemokines related to worse disease prognosis, such as IL-8, IL-6, CXCL1, and CCL20^75–77^, lowers the levels of anti-inflammatory markers^78^, and stimulates type I interferon production and expression of interferon-stimulated genes against Dengue virus infection^74^.

Considering that the direct interaction between celastrol and viral targets can mediate its antiviral activity^79,80^, we demonstrated its ability to bind M^pro^, RBD1, RDB2, and RdRp, which are essential to the viral life cycle ^19,24–27^. A previous study has described the celastrol interaction with the catalytic site of the SARS-CoV M^pro 81^. Docking simulations on the catalytic site of M^pro^ revealed that the celastrol binding energy was higher than that of most compounds tested here. Our models suggested that the high binding affinity of celastrol to M^pro^ involved the sigma Pi interaction between the E-ring O_2_ and His41, associated with the hydrogen bond between Thr25 and the hydroxyl group of A-ring C3, in agreement with literature reports^79,80^. In addition, our atomic distance analysis suggested that B-ring C6 interacted with Cys145 – a critical amino acid of M^pro^ catalytic site – by forming a Michael’s adduct; this inhibition mechanism of celastrol was also proposed in other molecular targets^82,83^. The anti-inflammatory activity of celastrol is associated with down-regulation of NF-kB pathway mediated by suppression of IKK activation (SETHI et al., 2006), probably due to the formation of Michael’s adduct between its quinone methide and Cys179 from IKK^80,84^. *In silico* studies have reported that other quinone derivatives exhibit antiviral properties associated with the ability to form Michael’s adduct^85^, such as the glucocorticoids methylprednisolone and dexamethasone used to treat severe Covid-19 patients^86,87^.

Next, we demonstrated that celastrol exerted antiviral and anti-inflammatory effect *in vitro*, which biologically validated our *in silico* findings here reported. Celastrol (250 to 1000 nM) reduced the viral progeny of SARS-CoV-2 in Vero CCL-81 and Vero CCL-81-ACE2 cells, as well as in the pulmonary and intestinal epithelial human cell lines Calu-3 and Caco-2, respectively. The anti-viral effect of celastrol is selective, since it does not inhibit influenza A virus replication (H1N1; PR8) in Madin-Darby Canine kidney cell line (MDCK) when tested at 150 to 600 nM^88^, but it significantly reduces HIV replication in U937 cells at 150 nM^89^, as we found for SARS-CoV-2.

As expected, the reuse of transcriptome data from SARS-CoV-2-infected NHBE cells, a normal human bronchial epithelial cell line, evidenced enriched biological pathways associated with non-hemostatic inflammation, including upregulation of the IL-6 gene expression. A recent review paper from our team has reported the relationship between serum IL-6 levels and different outcomes of Covid-19 patients^90^: compared with severe patients, critical and mild patients have higher and lower IL-6 levels, respectively; and non-survivors exhibit significantly higher IL-6 levels than survivors with Covid-19. *In vitro* assays using Calu-3, A549, and NHBE cells have confirmed that SARS-CoV-2 infection induces IL-6 production^30^. As IL-6 production is increased in both Covid-19 patients and *in vitro* SARS-CoV-2 infection models, we examined whether celastrol altered the levels of this cytokine *in vitro*. Celastrol at 500 and 1000 nM down-regulated the infection-induced IL-6 production in Caco-2 and Calu-3 cells, and such effect was associated with the decreased SARS-CoV-2 load. The inhibition of IL-6 production by celastrol could be associated or not with the reduced viral load observed on infected cells. The modulatory effect of celastrol over IL-6 release has been described *in vitro* under sterile inflammatory conditions^91^. In addition, this triterpene down-regulates IL-6 secretion and gene expression in PC-3 prostate carcinoma cells, in a NF-κB-dependent manner^92^, suppresses LPS-induced IL-6 production in RAW264.7 macrophages^93^, and decreases the IL-6 concentration and mRNA expression but not the virus title and mRNA expression in influenza A-infected MDCK cells^88^. Of note, celastrol reduced lung injury and the release of proinflammatory mediators into the pulmonary airways, including IL-6 production, in an experimental model of acute respiratory distress syndrome (ARDS)^94^.

Our findings demonstrated that celastrol was able to reduce both SARS-CoV-2 viral load and IL-6 production in two human cell lines, suggesting that this compound exerts anti-SARS-CoV-2 effects of clinical relevance to face the current pandemic. Several pharmacological applications of celastrol are associated with Covid-19 severity, including comorbidities such as obesity, diabetes, hypertension, and metabolic syndrome^95–98^, and pathological events such as non-homeostatic production of pro-inflammatory cytokines, thrombus generation, and formation of neutrophil extracellular traps^99–101^. Celastrol is currently under clinical trials to treat a variety of diseases, including different types of cancer, neurodegenerative disorders, and inflammatory conditions such as rheumatoid arthritis, psoriasis, and Crohn’s disease^102–106^.

Despite its potential therapeutic effects, there are still some limitations to the use of celastrol, such as its low solubility that results in poor bioavailability, *in vitro* and *in vivo* toxicity, and adverse effects that remain to be evaluated^107,108^. Celastrol was not cytotoxic to all cell lines tested herein, even at the highest concentration (1000 nM), corroborating other reports using Caco-2 cells^109^. At 1000 nM, this compound reduced the SARS-CoV-2-cytopathic effect on Vero CCL-81 cells. A recent study using a mouse model of acute toxicity has demonstrated that celastrol is safe even when orally administered at a high dose (62.5 mg/kg body weight)^110^. Intraperitoneal administration of celastrol (0.25 mg/kg body weight) increases the gamma irradiated-mice survival rate by around 70^78^. To surpass celastrol toxicity, solubility, and pharmacokinetic issues, several pharmaceutical approaches have been proposed, such as nanoencapsulation, liposomes, and sugar-silica nanoparticles^111–113^. Celastrol has been considered a lead drug for several human illnesses, but its toxicity to humans remains to be determined^64,73,114^.

To the best of our knowledge, the present study is the pioneer to use the combination between bioinformatic tools and biological approaches to demonstrate that celastrol inhibits the SARS-CoV-2 replication in non-human and human cell lines, and down-regulates IL-6 secretion from infected-human cell lines, reinforcing that celastrol is a potential repurposed drug to treat Covid-19.

## 5. Acknowledgements

We are thankful to the Brazilian National Council for Scientific and Technological Development (CNPq, grant #312606/2019-2 to M.D.B.), the Coordination for the Improvement of Higher Educational Personnel (CAPES, Finance Code 001), and the São Paulo Research Foundation (FAPESP, grant #20/05270-0 to V.L.D.B.).

## 6. Competing Interests

The authors declare no competing interests.

## Notes

### Competing Interest Statement

The authors have declared no competing interest.

