## Supplementary Figure S1 for "Drug repurposing to face Covid-19: Celastrol, a potential leading drug capable of inhibiting SARS-CoV-2 replication and induced inflammation"

**Figure S1.** Fuzo *et al.*

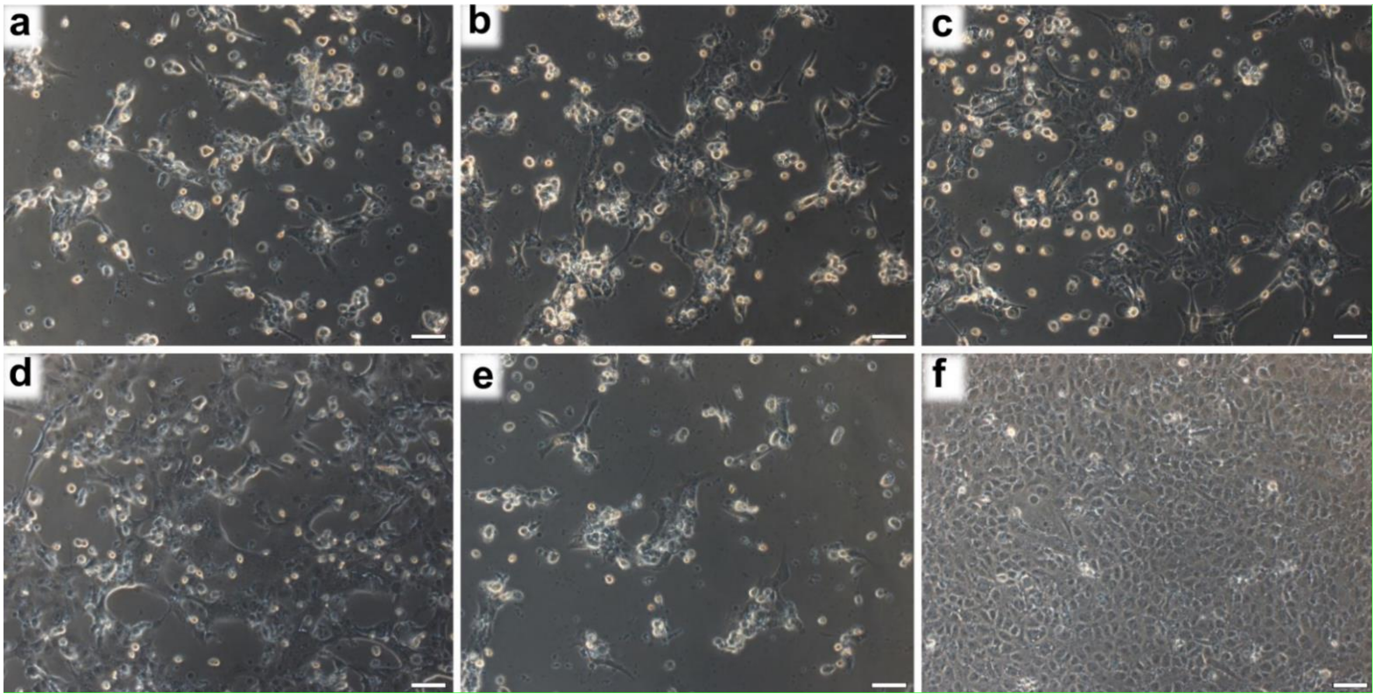


**Figure S1**. **Celastrol reduces SARS-CoV-2 cytopathic action.** Optical microscopy analysis of Vero CCL-81 cells infected with SARS-CoV-2 (MOI = 1.0) and treated with celastrol at the concentrations of (**a**) 125, (**b**) 250, (**c**) 500, and (**d**) 1000 nM, for 48 hours. Infected cells not treated with celastrol were used as the positive control (**e**). Cells incubated only with culture media were used as the negative control (**f**). Bar = 100 μm.
